# Deep speech-to-text models capture the neural basis of spontaneous speech in everyday conversations

**DOI:** 10.1101/2023.06.26.546557

**Authors:** Ariel Goldstein, Haocheng Wang, Leonard Niekerken, Zaid Zada, Bobbi Aubrey, Tom Sheffer, Samuel A. Nastase, Harshvardhan Gazula, Mariano Schain, Aditi Singh, Aditi Rao, Gina Choe, Catherine Kim, Werner Doyle, Daniel Friedman, Sasha Devore, Patricia Dugan, Avinatan Hassidim, Michael Brenner, Yossi Matias, Orrin Devinsky, Adeen Flinker, Uri Hasson

## Abstract

Humans effortlessly use the continuous acoustics of speech to communicate rich linguistic meaning during everyday conversations. In this study, we leverage 100 hours (half a million words) of spontaneous open-ended conversations and concurrent high-quality neural activity recorded using electrocorticography (ECoG) to decipher the neural basis of real-world speech production and comprehension. Employing a deep multimodal speech-to-text model named Whisper, we develop encoding models capable of accurately predicting neural responses to both acoustic and semantic aspects of speech. Our encoding models achieved high accuracy in predicting neural responses in hundreds of thousands of words across many hours of left-out recordings. We uncover a distributed cortical hierarchy for speech and language processing, with sensory and motor regions encoding acoustic features of speech and higher-level language areas encoding syntactic and semantic information. Many electrodes—including those in both perceptual and motor areas—display mixed selectivity for both speech and linguistic features. Notably, our encoding model reveals a temporal progression from language-to-speech encoding before word onset during speech production and from speech-to-language encoding following word articulation during speech comprehension. This study offers a comprehensive account of the unfolding neural responses during fully natural, unbounded daily conversations. By leveraging a multimodal deep speech recognition model, we highlight the power of deep learning for unraveling the neural mechanisms of language processing in real-world contexts.

## Introduction

One of the ultimate goals of our collective research endeavor in human neuroscience is to model and understand how the brain supports dynamic, context-dependent behaviors in the real world. Perhaps the most distinctly human behavior—and the focus of this paper—is our capacity for using language to communicate our thoughts to others during free, open-ended conversations. Natural language, as it occurs in daily conversations, is highly complex. It encompasses numerous linguistic rules, sub-rules, and exceptions and is influenced by discourse context, meaning, dialect, and other factors ^1–3^. Traditionally, neurolinguistics has approached real-world language’s complex and multidimensional nature by employing an incremental divide-and-conquer strategy. Individual labs have employed clever experimental manipulations to isolate and computationally model specific aspects of language processing divorced from the broader context. The implicit aspiration behind this collective effort is to eventually integrate these fragmented studies into a comprehensive neurocomputational model of natural language processing ^4–6^. After decades of research, however, there is increasing awareness of the gap between controlled laboratory experiments and the complexity of everyday life ^7,8^. Models and theories developed in a particular experimental context often fail to generalize to other, more ecological contexts ^9–11^. To make matters even more challenging, language and communication are spontaneous, dynamic, and fundamentally contextual, unamenable to many core tenets of experimental design (e.g., repetition and trial averaging ^12^.

To study the neural basis of natural language processing in the real world, we developed a new electrocorticography (ECoG) paradigm to measure human neural activity in real-world, naturalistic contexts at scale—during hundreds of hours of open-ended conversations, comprising both speech production and speech comprehension. Unlike traditional ECoG studies, which typically rely on controlled experiments performed over short durations, our dense-sampling paradigm enabled continuous 24/7 recording of ECoG and speech data during extended days-to week-long stays in the hospital epilepsy unit. This ambitious effort resulted in a uniquely large ECoG dataset of natural conversations: four patients recorded during free conversations, yielding approximately 50 hours (226,776 words) of neural recordings during speech comprehension and 50 hours (283,736 words) during speech production in real-world settings. Modeling the neural basis for natural language processing in open-ended conversations presents an unprecedented challenge, given that we have no experimental control and no two conversations are the same. Patients are free to say whatever they want, whenever they want, with no experimental intervention. Some conversations are with family members and friends, some with the hospital support team, and some with the doctors. Each conversation has its own context and purpose.

To model neural activity during real-life conversations, we relied on a new family of deep language models that can accommodate the complexity, multidimensionality, and context-dependent nature of language processing ^13–16^. However, most modern language models are trained to process large corpora of text; processing continuous speech into discrete words poses a cognitive challenge for these models, as it requires continuous parsing of speech sounds into meaningful lexical units. In this work, we leverage a multimodal neural network model called Whisper that learns to process spoken language in real-world contexts ^17^. The Whisper architecture incorporates both a multilayer “encoder” network and a multilayer “decoder” network (Fig. 1): the “encoder” maps continuous speech inputs into a high-dimensional embedding space capturing acoustic features of *speech*; the decoder maps discrete lexical (i.e., word) tokens into an embedding space capturing the contextual structure of *language*, similar to other autoregressive language models ^18–20^. We extracted two types of embeddings from Whisper for each word heard or produced by the patients in each conversation: (1) embeddings extracted from the “encoder” network, which we refer to as “speech embeddings”; and (2) embeddings extracted from the “decoder” network, which we refer to as “language embeddings”.

**Figure 1.**
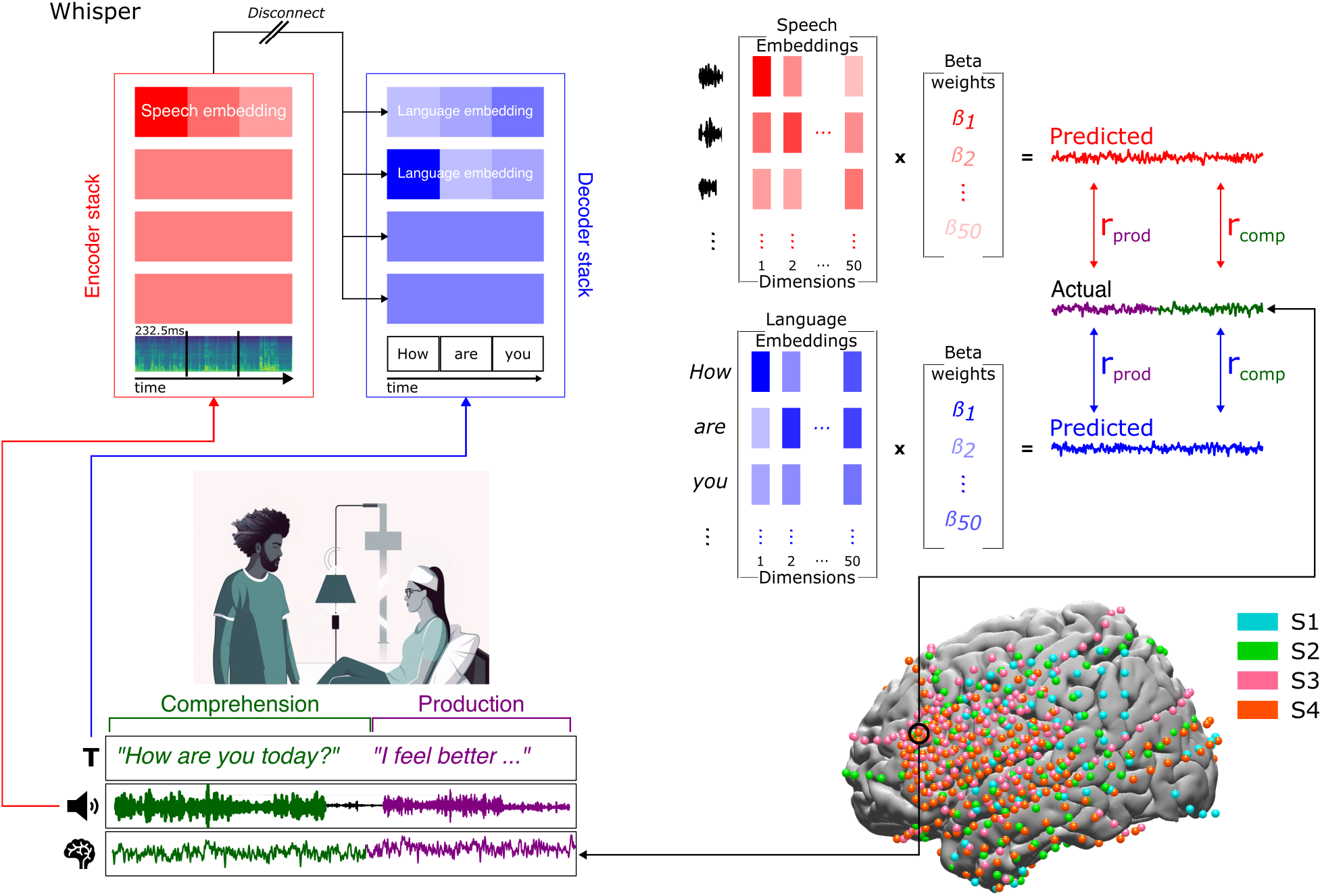
An ecological, dense-sampling paradigm for modeling neural activity during real-world conversations. Continuous brain signal monitoring of four ECoG patients during their interactions with hospital staff, family, and friends provides a unique opportunity to investigate real-world social communication. Simultaneously recorded verbal interactions are transcribed and segmented into production (*purple*) and comprehension (*green*) components **(bottom left)**. The Whisper model was employed to process the audio recordings and corresponding transcriptions. For each word, we extracted “speech embeddings” from Whisper’s encoder network (red) and “language embeddings” from Whisper’s decoder network (blue) **(top left)**. The embeddings were reduced to 50 dimensions using PCA. Linear regression predicted neural signals from the speech embeddings (*red*) and language embeddings (*blue*) across tens of thousands of words. To evaluate encoding model performance, we calculated the correlation between predicted and actual neural signals for left-out test words. This process was repeated for each electrode and each lag, using a 25 ms sliding window ranging from –2000 to +2000 ms relative to word onset **(top right)**. Brain coverage across four participants comprising 654 left hemisphere electrodes (**bottom right**).

Prior studies have demonstrated that embeddings in acoustic encoders like wav2vec learn the phonemic structure of natural speech^21^, while embeddings in language models like GPT-2 ^20,22^ learn the syntactic and semantic structure of natural language ^15,23–25^. Both of these types of models have begun to show promising results in modeling neural responses, in low-level auditory cortex and higher-level language areas, respectively ^26–33^. The multimodal architecture of Whisper enables us to jointly model and dissociate speech and language processing using components of the same model. Furthermore, capitalizing on the high spatiotemporal resolution of ECoG, Whisper allows us to capture the temporal interplay between speech comprehension and spontaneous speech production in completely unconstrained real-world conversations.

We constructed two sets of electrode-wise encoding models to estimate a linear mapping from both speech embeddings and language embeddings extracted from Whisper to the neural activity for each word during both speech production and comprehension. Our encoding moles revealed a remarkable alignment between the human brain and the internal representations of the speech-to-text model. Speech embeddings better captured cortical activity in both lower-level speech perception and speech production areas, including the superior temporal cortex and precentral gyrus. Linguistic embeddings, on the other hand, were better aligned with higher-order language areas like the inferior frontal gyrus and angular gyrus. Additionally, prior to each word onset during speech production, we observed a temporal sequence from language-to-speech encoding across cortical areas; during speech comprehension, we observed the reverse progression from speech-to-language encoding after word articulation. Our findings provide strong evidence that deep speech-to-text models can provide a novel modeling framework for the neural basis of both language production and comprehension across large volumes of real-world data without sacrificing the diversity and richness of natural language.

## Results

We collected continuous 24/7 recordings of ECoG and speech signals from four patients as they engaged in spontaneous conversations with their family, friends, doctors, and hospital staff during their entire days-long stay at the epileptic unit. Across the four patients, we recorded neural signals from 676 intracranial electrodes (Fig. 1). Because only one of the four patients had 22 electrodes implanted in the right hemisphere, we focused on left hemisphere electrodes (n = 654) in our analyses; 10 electrodes were excluded due to incomplete recordings, leaving 644 electrodes for analysis. We obtained extensive coverage of key language areas, including in the inferior frontal gyrus (IFG, also known as Broca’s area; n = 75) and superior temporal gyrus (STG; n = 45), with a sparser sampling of angular gyrus (AG; n = 35). We built a preprocessing pipeline to identify the occurrence of speech, remove identifying information, transcribe each conversation, and align each word with the concurrent ECoG signals. We then divided the data into two categories: comprehension (when patients were listening to speech) and production (when patients were producing speech). This unconstrained recording paradigm yielded neural activity from multiple electrodes (102–230 electrodes) for dozens of hours (17–37 hours), comprising tens of thousands of words (79,654–213,473 words) per patient. A comprehensive description of the speech collected, patient demographics and clinical characteristics see Table S1. Preprocessing procedures can be found in the Materials and Methods section.

In our dataset, each conversation is unique, allowing patients to express themselves freely without any intervention from experimenters. To accommodate the complex and multimodal nature of real-world speech, we employed Whisper, a multimodal speech-to-text model that can learn to align real-world speech recordings with high-level linguistic meaning with human-level precision ^17^. We input the acoustic speech recordings and manually-transcribed text from our conversations to Whisper. To leverage the multimodal architecture of Whisper, we separately extract both “speech embeddings” and “language embeddings” for each word in every conversation (Fig. 1; Materials and Methods): speech embeddings were extracted from the encoder network based on continuous speech inputs; language embeddings were extracted from the decoder network. During the extraction of embeddings, we disconnect the cross attention in the Whisper model and separate it into a speech encoder stack and a language decoder stack.

What features of real-world conversations do these speech and language embeddings encode? To visualize the information encoded in these embeddings, we used t-SNE to project the multidimensional embeddings (3840-d for speech and 384-d for language, sampled from each layer of the encoder and decoder) onto two-dimensional manifolds (Fig. 2 & S1). We found that speech embeddings are clustered according to known acoustic features of speech, such as phonemes (Fig. 2A), but not lexical information, such as part of speech (PoS; Fig. 2C). On the other hand, the language embeddings are clustered according to part of speech (Fig. 2B) and, to a lesser extent, phonemes (Fig. 2D). Similar clustering results were obtained for t-SNE projections of manner of articulation (MoA) and place of articulation (PoA), see supplementary Fig. S1A-D. To quantify which features are encoded in these embeddings, we assessed classification performance at each layer for four different types of features: phoneme categories (27 classes), place of articulation (PoA, 9 classes), manner of articulation (MoA, 9 classes), and part of speech (5 classes) ^34^. The speech embeddings yielded robust classification performance for all features (Figs. 2E & S1E), with a particularly strong classification of phonemes at the last layer of the encoder (accuracy = 0.54, chance level = 0.04, 27 classes), PoA (accuracy = 0.48, chance level = 0.11, 9 classes), and MoA (accuracy = 0.63, chance level = 0.11, 9 classes). The final (fourth) layer of the Whisper encoder network yielded the best classification of phonemes, PoA, and MoA (Fig. S1E), so we extracted speech embeddings from this layer for further analysis. The language embeddings, on the other hand, yielded a stronger classification of PoS (accuracy = 0.66, chance level= 0.20, 5 classes). Classification of PoS was highest for the final layers of the Whisper decoder network (Fig. S1F), so extracted language embeddings for the second-to-last (third) layer, in keeping with prior work showing that late-intermediate layers of autoregressive language models provide the best predictions for neural activity ^29,31,35^. Similar results were obtained for the top language decoder layer.

**Figure 2.**
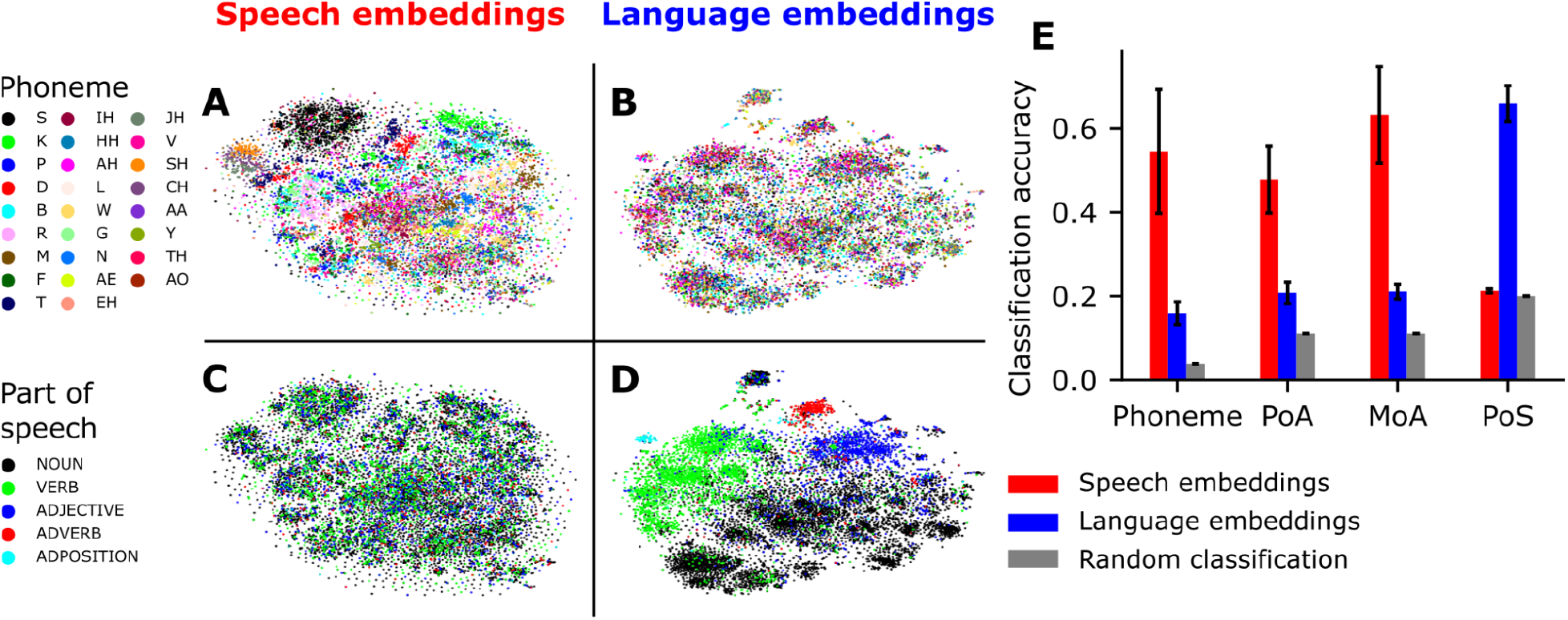
Representations of phonetic and lexical information in Whisper. (A-D) Speech embeddings and language embeddings were visualized in a two-dimensional space using t-SNE. Each data point corresponds to the embedding for either an audio segment (speech embeddings from the encoder network) or a word token (language embeddings from the decoder network) for a unique word (averaged across all instances of a given word). Clustering according to phonetic categories is visible in speech embeddings (A) but far less prominent in language embeddings (B). Clustering according to lexical information (part of speech) is visible in language embeddings (D) but not in speech embeddings (C). **(E)** Classification of phonetic and lexical categories based on speech and language embeddings. We observed robust classification for phonetic information based on speech embeddings. We also observed robust classification for parts of speech based on language embeddings.

Together, these findings provide evidence that Whisper’s speech embeddings implicitly encode acoustic features of natural speech, while the language embeddings implicitly encode lexical properties. Note that Whisper was trained end-to-end to predict upcoming words given the audio as input; the encoder was not explicitly trained to recognize phonemes or manner of articulation, and similarly, the decoder was not trained to recognize parts of speech. The emergent representation of these psycholinguistic features in different architectural components of the model motivates our use of Whisper’s speech and language embeddings to model speech- and language-related features of human brain activity.

To assess whether the embeddings extracted from Whisper can capture neural activity during natural conversations, we constructed two sets of encoding models based on speech embeddings and language embeddings during both speech production and speech comprehension. The encoding models estimate a linear mapping between the Whisper embeddings and neural activity for each word in the training set. Subsequently, we used the trained encoding models to predict the neural activity for each word at each electrode in left-out conversations (Fig. 1). A separate encoding model was trained for each electrode at various time points, ranging from -2000 ms to 2000 ms relative to the word onset (time 0). The performance of the encoding model was evaluated by calculating the correlation between the predicted and actual neural signals for the held-out conversations using ten-fold cross-validation. All analyses were adjusted for multiple comparisons using a non-parametric procedure to control the family-wise error rate (FWER).

Whisper’s speech and language embeddings predicted neural activity with remarkable accuracy across conversations comprising hundreds of thousands of words, during both speech production and comprehension, for numerous electrodes in various regions of the cortical language network (Fig. 3). These brain regions are known to be involved in auditory speech processing (e.g., superior temporal gyrus; STG), language comprehension and production (e.g., inferior frontal gyrus, IFG), somatomotor (SM) planning and execution (e.g., precentral and postcentral gyrus; preCG, postCG), and high-level semantic cognition (e.g., angular gyrus and temporal pole; AG, TP) ^36,37^. Speech embeddings yielded more significant electrodes than language embeddings for both production (274 versus 154, chi-square (1, *N* = 644) = 49.55, *p* < 0.001) and comprehension (186 versus 135, chi-square (1, *N* = 644) = 4.09, *p* < 0.05). Remarkably, the predicted signals were strongly correlated with the actual signals—up to 0.5 Pearson correlation across lags—across hours of left-out speech segments. Similar results were obtained when using the last layer of the decoder stack or the unimodal GPT-2 (Fig. S2).

**Figure 3.**
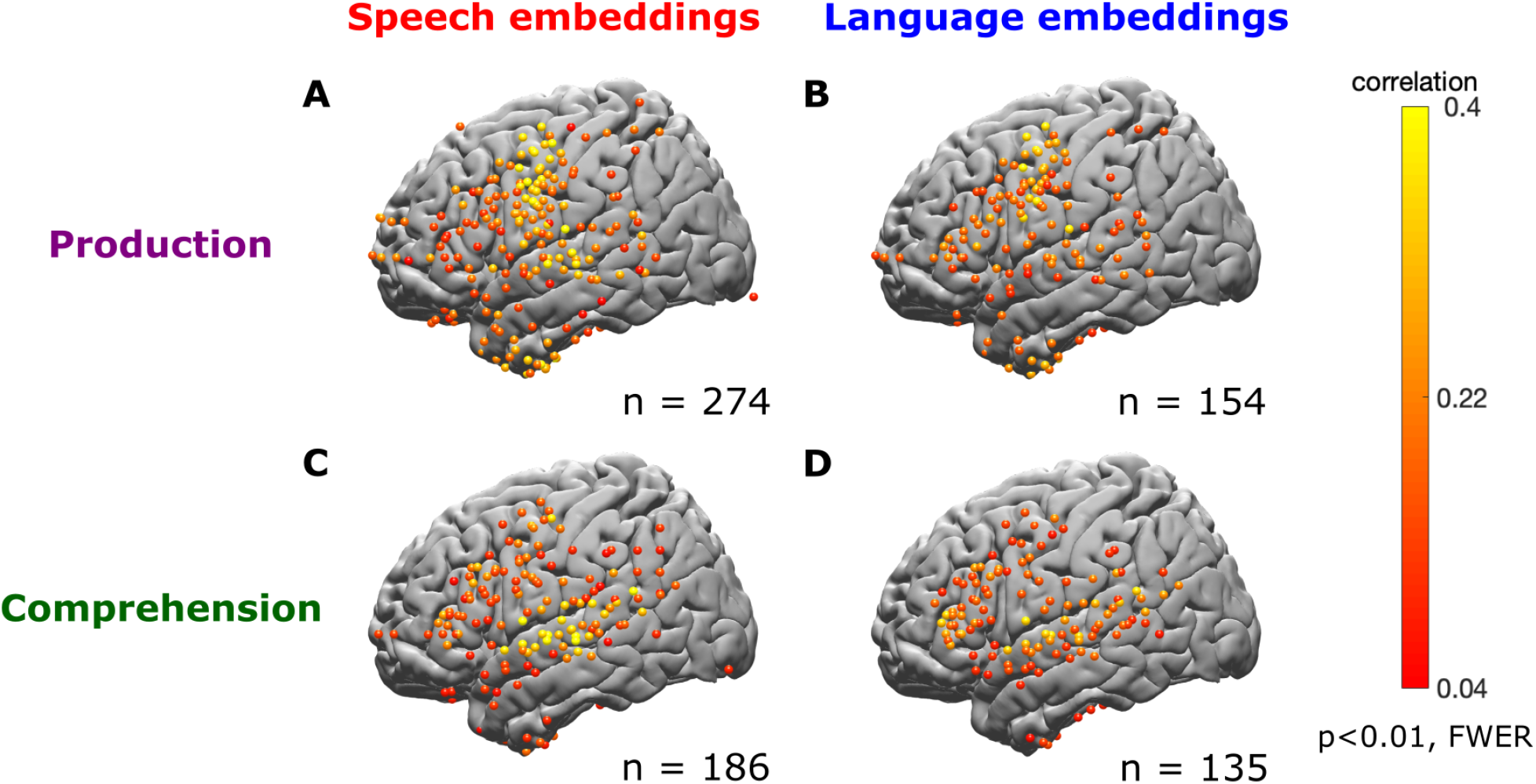
Speech and language encoding model performance during speech production and comprehension. Encoding performance (correlation between predicted and actual neural activity) for each electrode for speech (acoustic) and language (lexical) models during speech comprehension (∼50 hours, 289,971 words) and speech production (∼50 hours, 230,238 words). The plots illustrate the correlation values associated with the encoding for each electrode, with the color indicating the highest correlation value (across lags). **(A)** Modeling speech production using speech embeddings revealed significant electrodes in somatomotor areas (SM) and superior temporal gyrus (STG), as well as the inferior frontal gyrus (IFG; Broca’s area), temporal pole (TP), angular gyrus (AG), and posterior middle temporal gyrus (pMTG; Wernicke’s area). **(B)** Employing language embeddings for speech production also highlighted similar regions but notably fewer electrodes (with lower correlations) in SM and STG and higher correlations in IFG. **(C)** Modeling speech comprehension using speech embeddings resulted in significant electrodes in auditory and language areas, particularly the STG, TP, IFG, and Wernicke’s areas. **(D)** Similarly, when using language embeddings for speech comprehension, a comparable pattern emerged with slightly lower performance in STG and higher in the IFG. Overall, the findings demonstrate specific cortical regions associated with speech production and comprehension, showcasing the distinct contributions of speech and language embeddings in decoding neural activity.

We could accurately predict speech and language-related neural activity during new conversations in numerous individual electrodes (Fig. 4). During spontaneous speech production (Fig. 4A), we observed organized hierarchical processing, where articulatory areas along the preCG and postCG, as well as STG, better correlated with speech embeddings (red), while higher-level language areas such as IFG, pMTG, and AG better correlated with language embeddings (blue). A similar hierarchical organization was evident in speech comprehension (Fig. 4B): perceptual areas such as STG and somatomotor areas like preCG and postCG showed a preference for speech embeddings, while higher-level language areas, including IFG and AG, displayed a preference for language embeddings. Intriguingly, we observed a pronounced preference for speech information in the TP during speech production (Figs. 4, S3). Our predictions had a high level of precision, with a correlation between predicted and actual neural responses ranging from 0.2 to 0.5 across electrodes and models. We replicated these electrode-level findings at the level of anatomically-defined regions of interest (averaging encoding performance across electrodes at each ROI; Fig. S3). Similar results were obtained when using the language embeddings without detaching them from the encoder network (Fig. S4). These high correlations were achieved for hundreds of thousands of words and tens of hours of speech from previously unseen, unique conversations not used to train the encoding model (Fig. 4).

**Figure 4.**
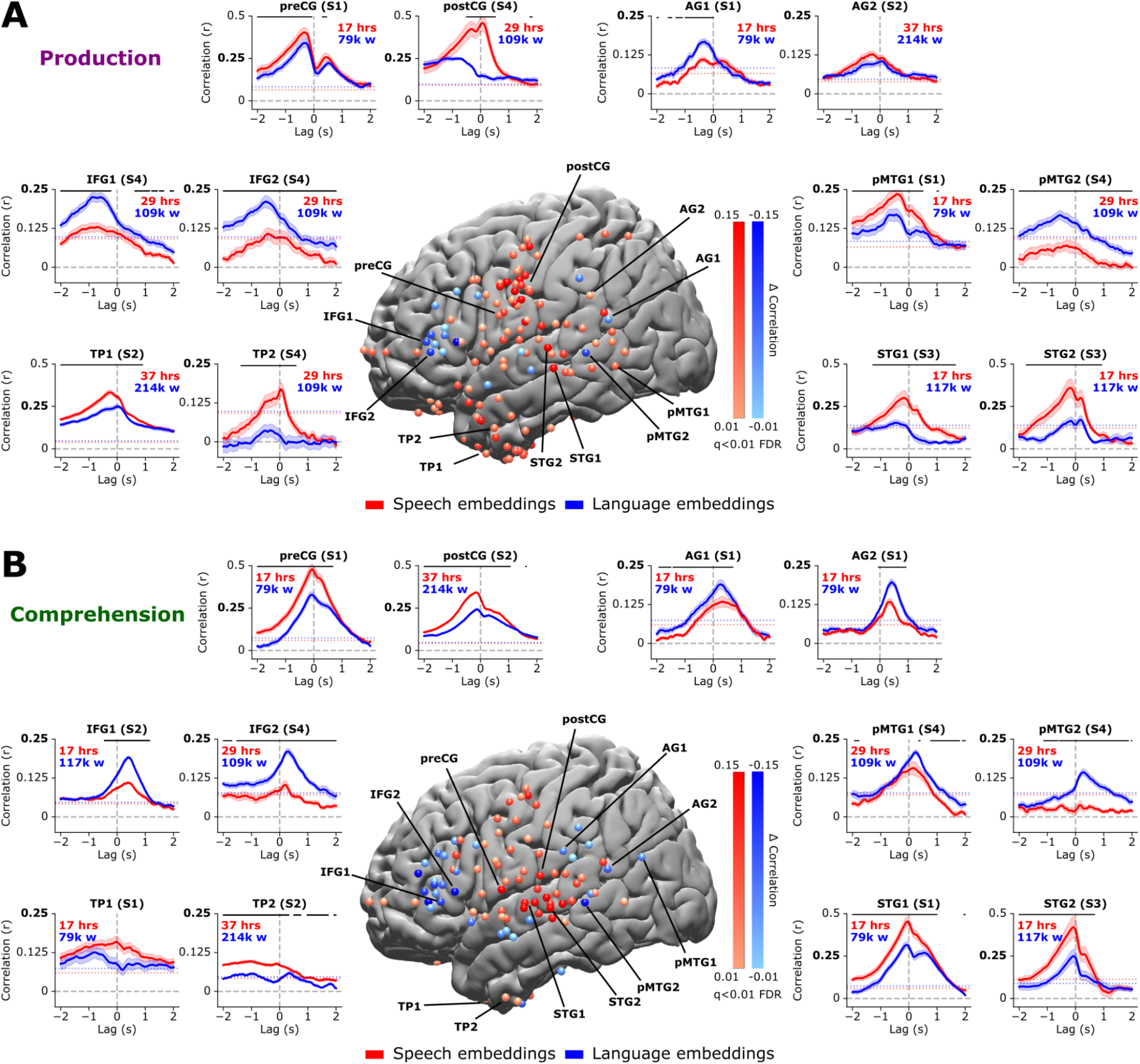
Mixed selectivity for speech and language embeddings during both speech production and comprehension. (A) Contrast map of electrodes exhibiting significantly higher encoding for speech embeddings (red) and significantly higher encoding for language embeddings (blue). Surrounding plots display encoding performance during speech production for selected individual electrodes across different brain areas and patients. Models were estimated separately for each lag (relative to word onset at 0 s) and evaluated by computing the correlation between predicted and actual neural activity. The dashed horizontal line indicates the statistical threshold (*q* < 0.01, FDR corrected). The speech encoding model (red) achieved correlations of up to 0.5 when predicting neural responses to each word over hours of recordings in the superior temporal gyrus (STG), precentral gyrus (preCG), and postcentral gyrus (postCG). The language encoding model yielded significant accuracies (correlations up to 0.25) and outperformed the speech model in IFG and AG. **(B)** Contrast map of speech embeddings (red) versus language embeddings (blue) during speech comprehension. Matching the flow of information during conversations, encoding models accurately predicted neural activity ∼500 ms before word onset during speech production and 300 ms after word onset during speech comprehension.

Evaluating encoding models at each lag relative to word onset allows us to trace the temporal flow of linguistic information across speech-related ROIs during the production and comprehension of natural conversations. First, in congruence with the flow of information during speech production, language encoding in IFG peaked first around 500 ms before word onset (M = -480 ms, SD = 284 ms), whereas, in somatomotor areas (SM; comprising preCG and postCG), the speech model encoding peaked significantly closer to speech onset (M = -305 ms, SD = 391 ms, t(80) =1.95, p < 0.05; Fig. 5A). Interestingly, the temporal dynamics of speech encoding in the SM and auditory speech areas along the STG were aligned. This may indicate an efferent copy between motor and auditory areas during speech production ^38–40^. A reverse dynamic was observed during speech comprehension. During speech comprehension, speech areas along the STG peaked shortly after word-onset (M = 55 ms, SD = 189 ms), while language model encoding in IFG peaked significantly later, around 300 ms after word onset (M = 292ms, SD = 73ms, t(60) = -6.35, p < 0.001; Figs. 5B, S5). During speech comprehension, encoding in motor areas exhibited higher variability and was less interpretable, as shown in Fig. 5B. Finally, we found an unexpected temporal pattern of speech encoding during speech production: peak encoding performance proceeded from dorsal SM to middle SM and finally to ventral SM prior to word articulation (Fig. 5C).

**Figure 5.**
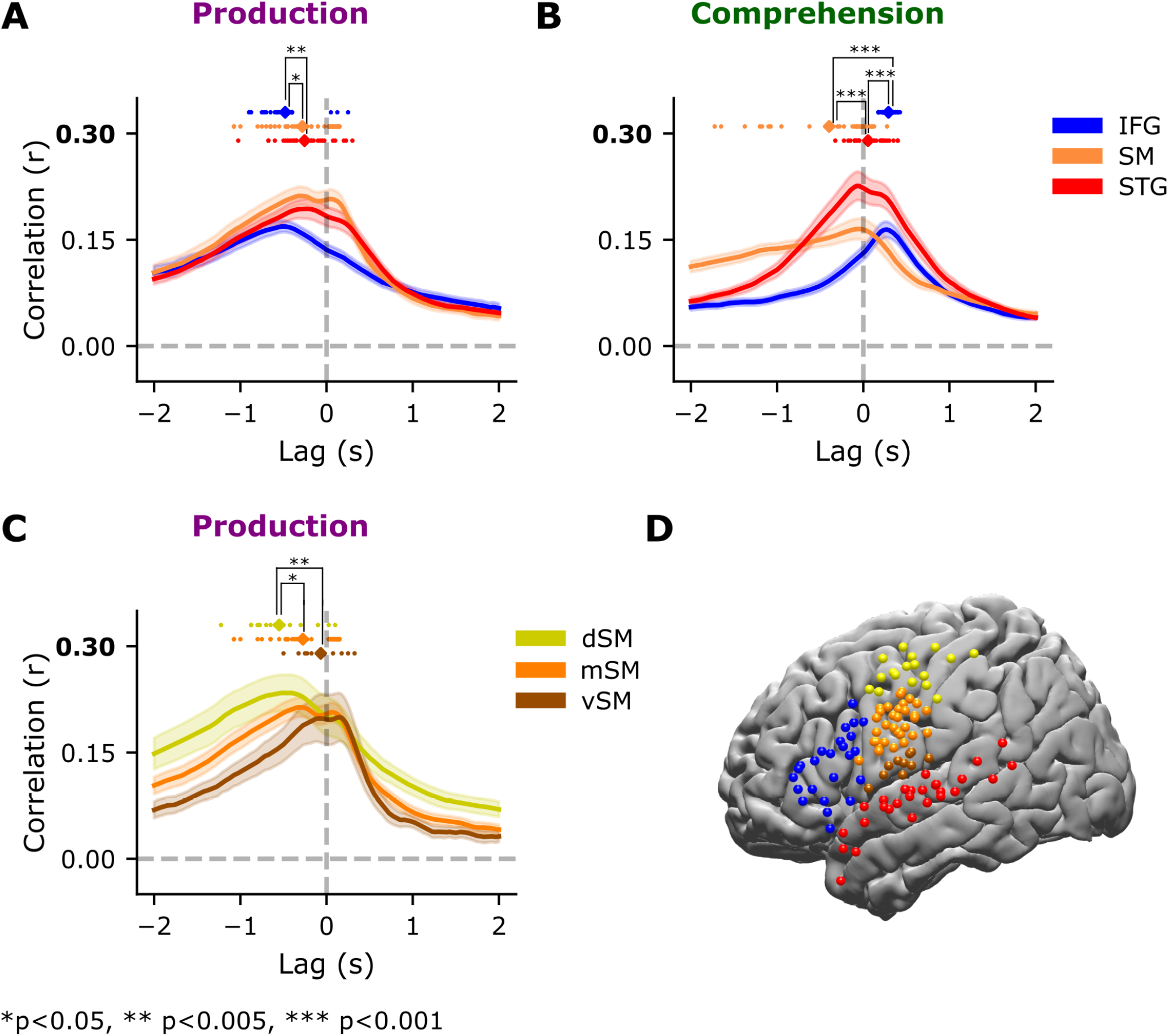
Temporal dynamics of speech production and speech comprehension across different brain areas. Based on tuning preferences for each ROI, we assessed temporal dynamics using the language model for IFG and the speech model for STG and SM. Colored dots show the lag of the encoding peak for each electrode per ROI. To determine significance, we performed two-sample permutation tests between encoding peaks. **(A)** During speech production, encoding for language embeddings in IFG peaked significantly before speech embeddings in SM and STG. **(B)** The reverse pattern was observed during speech comprehension: encoding performance for language embeddings encoding in IFG peaked significantly after speech encoding in SM and STG. **(C)** For speech production, we observed a temporal pattern of encoding peaks shifting toward word onset within SM, proceeding from dorsal (dSM) to the middle (mSM) to ventral (vSM). **(D)** Map showing the distribution of electrodes per ROI.

Lastly, we mapped the electrode-wise selectivity for production versus comprehension across the cortical language network using embeddings for either speech embeddings (Figs. 6A, S6) or language embeddings (Figs. 6B, S6). In comparing production versus comprehension, we observed a different hierarchical organization, where speech areas in STG and language areas in anterior and medial IFG yielded higher encoding performance during speech comprehension (Fig. 6AB, green), while posterior IFG and SM (preCG and postCG), as well as the temporal pole (TP), yielded higher encoding performance during speech production (Fig. 6AB, purple). Similar results were seen for language embeddings (Fig. 6B). Interestingly, these results suggest a gradient from speech comprehension at the anterior part of IFG to speech production at posterior IFG toward SM areas. We found that SM areas play a surprisingly large role in real-life unconstrained conversations in terms of both speech and language features.

**Figure 6.**
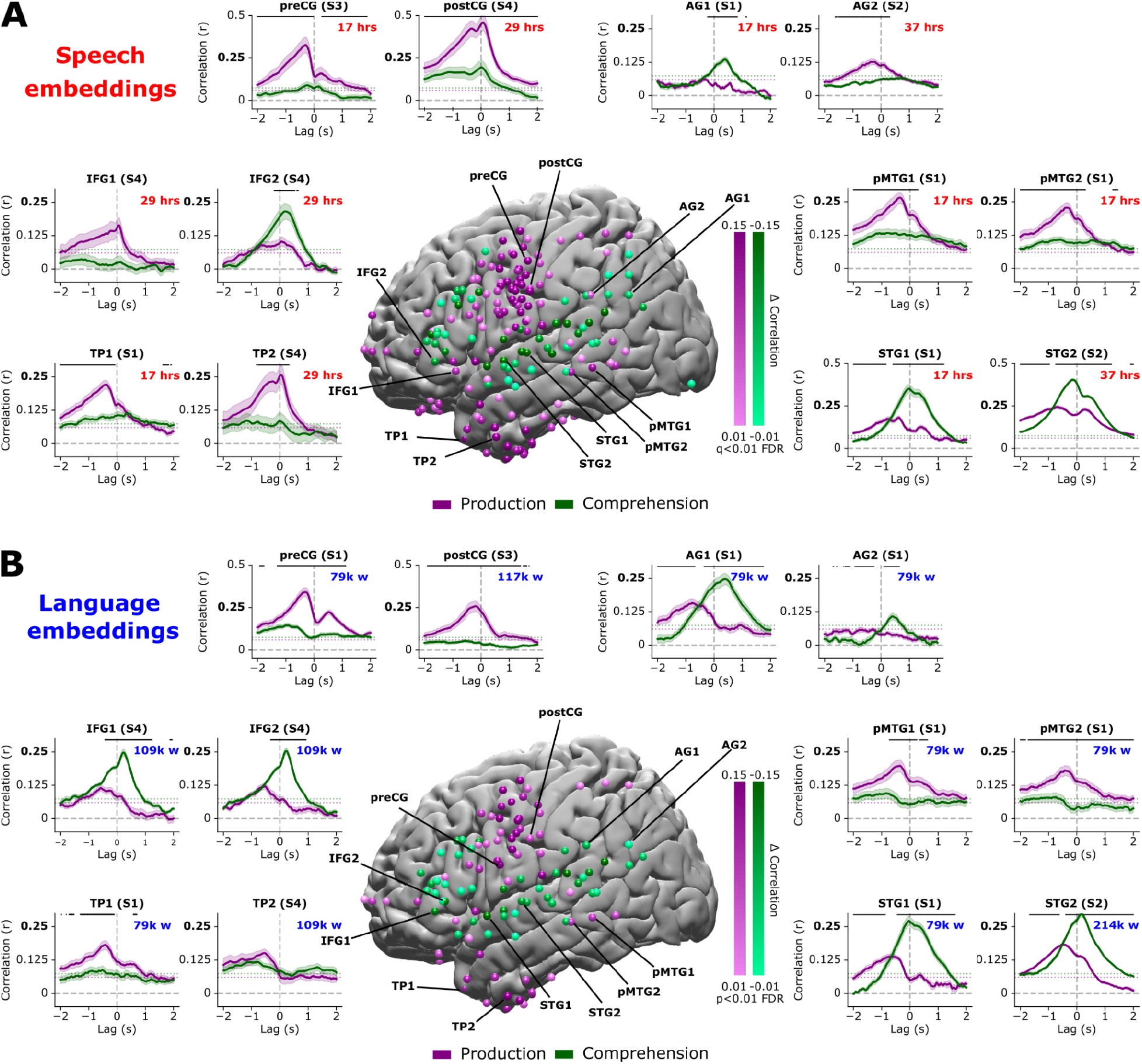
Mixed selectivity for speech production and comprehension. **(A)** Contrast map of the electrodes exhibiting significantly higher encoding during speech production (purple) and significantly higher encoding during speech comprehension (green) using encoding models based on speech embeddings. Surrounding plots display encoding performance during speech production and speech comprehension for selected electrodes across different brain areas and patients. Models were estimated separately for each lag (relative to word onset at 0 s) and evaluated by computing the correlation between predicted and actual neural activity. The dashed horizontal line indicates the statistical threshold (*q* < 0.01, FDR corrected). STG, IFG, and AG exhibited higher encoding during speech comprehension (green). The precentral gyrus (preCG), postcentral gyrus (postCG), medial temporal gyrus (MTG), and temporal pole exhibited higher encoding during speech production (purple). **(B)** Contrast map of encoding performance during speech production (purple) versus speech comprehension (green) based on language embeddings.

## Discussion

In this study we modeled neural activity in humans as they freely speak to and listen to others in interactive, real-world conversations. We introduce a dense-sampling paradigm for modeling the neural machinery driving everyday conversations in a way that embraces their complex and contextual nature. For the first time, we recorded one hundred hours—comprising half a million words—of spontaneous open-ended conversations concurrently with high spatiotemporal resolution neural activity using intracranial recordings. The unprecedented size of this dataset provides us with a detailed look at the richness of human conversations as they unfold in real-world contexts. Using speech and language embeddings from a multimodal speech-to-text model, we could predict neural activity with remarkable precision in both lower-level perceptual and motor, as well as higher-level language areas, across hundreds of thousands of words in free conversations. The model’s generalization performance across many hours of everyday conversations implies that we can now predict neural responses in language areas for most conversations without simplifying or controlling speech.

Our encoding models revealed a distributed processing hierarchy in which sensory areas along the superior temporal gyrus (STG) and somatomotor areas (SM) along the precentral gyrus (preCG) were better modeled by speech embeddings (red, Fig. 3), while higher-order language areas in the inferior frontal gyrus (IFG), as well as posterior temporal and parietal cortex, were better modeled by language embeddings (blue, Fig. 3). This was true for both speech production (Fig. 3A) and speech comprehension (Fig. 3B). These results recapitulate the known hierarchy of natural language processing ^41,42^. At the same time, electrode-wise selectivity for speech or linguistic information was mixed across most brain areas: higher-level linguistic information was encoded in speech perception and speech articulation areas (Fig. 4, individual electrodes, and Figs. S2, S3 & S6 for ROI level analysis) and speech information was encoded in higher-order language areas (Fig. 4, individual electrodes, and Figs. S2, S3 & S6 for ROI level analysis). This mixed selectivity is common in both biological and artificial learning systems that are “directly” fit to the complex structure of their inputs ^13^, and may support the high-dimensional neural representational space needed to encode the structure of natural language ^43^. Overall, these results are aligned with the known interaction between speech level and semantic level processing, where linguistic prediction can facilitate speech processing in auditory areas, and acoustic information can facilitate the processing of words in language areas ^44–48^.

Most of our knowledge about speech production relies on studies in which speakers are asked to articulate a fixed set of predetermined sentences ^49,50^. This study provides insights into the neural mechanisms underlying the production and articulation of spontaneous thoughts during open-ended unconstrained conversations, a cognitive process that has so far remained enigmatic. Generally, motor areas along the preCG are more central for speech production (purple, Figs. 6, S6). However, many electrodes exhibit mixed selectivity for speech production and speech comprehension. For example, comprehension-related areas in the STG encode speech (and higher-level linguistic) information before word onset during speech production ^51,52^. Conversely, we could predict the neural activity for perceived speech even in motor areas associated with speech production (Fig. 3). This indicates that speech production and comprehension may rely on a shared neural system ^53–55^. This was also true for high-order language areas.

The high spatiotemporal resolution of our neural recordings allows us to map the temporal dynamics of speech and language encoding across cortical areas during everyday conversations. In IFG, our encoding models accurately predicted neural activity for each word ∼500 ms before word onset during speech production and ∼300 ms after word onset during speech perception. The temporal encoding sequence across brain areas was reversed when speaking versus listening. For example, during speech production, we observed that peak language encoding in IFG preceded peak speech encoding in somatomotor areas (SM). In contrast, during speech comprehension, peak speech encoding along STG preceded peak language encoding in the IFG. Notably, during speech production, we found strong alignment to speech embeddings in both articulation areas in SM and perceptual areas in STG, suggesting a potential coupling between motor and perceptual processes during speech production ^39,55^.

How should we interpret the relationship between the internal representations of Whisper and the human brain when processing human speech? There are two potential options to consider. The first option is that our encoding model effectively learns the transformation between two distinct codes for processing natural language. This is significant because it positions deep language models as a powerful computational tool to study and predict how the brain processes everyday conversations. They enable us to map out brain areas sensitive to speech and linguistic information during both speaking and listening across hours of context-rich natural language. This breakthrough was instrumental in modeling our unique, fully unconstrained conversational dataset.

The second interpretation is that there are shared computational principles for natural language processing between deep language models and the human brain. This is a stronger claim that suggests that deep learning can serve as a neurally-inspired cognitive model for human brain function. In particular, these models have leveled a challenge against traditional rule-based symbolic linguistic models of language representation and processing ^16^. Some arguments support the stronger theoretical claim. First, our encoding models established that a simple linear mapping between the internal neural activity in Whisper and the human brain yields remarkably high prediction performance. This suggests the two internal representations may be more similar than initially anticipated. Second, recent research has discovered shared computational principles between unimodal deep language models (such as GPT2 or BERT) and the human brain during comprehension of previously-recorded spoken stories ^29,35,56^. Our investigation goes beyond these previous studies by introducing a multimodal speech-to-language model and examining both comprehension and production of spontaneous speech. Our findings support linguistic theories that highlight the role of usage-based statistical learning in language acquisition and downplay the importance of classical rule-based symbolic linguistic models.

## Materials and Methods

### Preprocessing the speech recordings

We developed a semi-automated pipeline for preprocessing the dataset. The pipeline can be broken down into four steps:

1. *De-identifying speech:* All conversations in a patient’s room were recorded using a high-quality microphone and stored locally. These audio recordings contain sensitive information about the patient’s medical history and private life. To comply with HIPAA’s data privacy and security provisions for safeguarding medical information, any identifiable information (e.g., names of people and places) were censored. Given the sensitivity of this phase, we employed a research specialist dedicated to the manual de-identification of recordings for each patient.
2. *Transcribing speech:* Although many speech-to-text transcription tools have been developed, extracting text from 24/7 noisy, multi-speaker audio recordings is challenging. To achieve the transcription quality necessary for our preliminary analyses, we used a human-in-the-loop annotation pipeline integrated with human transcribers from MTurk.
3. *Aligning text to speech:* Text transcripts (i.e., sequences of words) must be aligned with the audio recordings at the individual word level to provide an accurate time stamp for the production of each word. We used the Penn Forced Aligner ^57^, which yields timestamps with 20-millisecond precision, to generate rough word onsets and offsets. We further improved this automated forced alignment by manually verifying and adjusting each word’s onset and offset times.
4. *Aligning speech to neural activity:* To provide a precise mapping between neural activity and the conversational transcripts, we engineered one of the ECoG amplifiers to record the output of the microphones directly. The concurrent recordings of the audio and neural signals allowed us to align both signals at about 20-millisecond precision.

### Speech embedding extraction

To prepare audio recordings for subsequent processing by the speech model, we downsampled the audio recordings to 16 kHz. Since Whisper is trained on 30 s audio segments, audio recordings were fed to the model using a sliding window of 30 s. The Whisper encoder’s internal representations are not aligned to discrete word tokens (as in the decoder); instead, the encoder embeddings correspond to temporal segments of the original audio input. To account for this, we used an approximation to extract embeddings for each word. To understand this approach, it is necessary to give an overview of the model architecture: In the first step, each 30 s audio segment is transformed to a spectrogram of size 3000 * 80, where 3000 corresponds to the number of hidden states and 80 to the number of mel features. Each hidden state represents a temporal segment of 25 ms of the original audio input, with a stride of 10 ms. The spectrogram is then passed through two convolutional layers, resulting in 1500 hidden states of 384 embedding dimensions each. As a result of the convolutional operations, each hidden state represents a temporal segment of 62.5 ms of the original audio input, with a stride of 20 ms. The mean word duration in our dataset was around 235 ms (M = 235 ms, SD = 169 ms). To extract embeddings on the word level, we concatenated the last ten hidden states, corresponding to 232.5 ms of the audio input. To temporally align the concatenated ‘word embedding’ to the word onset, we defined the endpoint of each sliding window to the word’s onset plus 232.5 ms. Thus, the extracted ‘word embedding’ does not contain any information prior to word onset after the spectrogram and convolution layers. Since our classification analysis indicated that embeddings extracted from the fourth encoder layer have the most structured representation of phonetic categories compared to embeddings extracted from other encoder layers (Fig. S1E), encoding analyses were performed using embeddings from the fourth encoder layer.

### Language embedding extraction

For each word, text transcripts corresponding to the 30 s context window were tokenized and given as contextual input to the decoder (M = 70 words in, SD = 28 words in a 30 s window). We extracted the embedding corresponding to the last word in the sequence. In line with previous results indicating that late-intermediate layers of language models show the best encoding performance for neural data ^58^, we extracted embeddings from the third decoder layer.

### Visualization of embedding space

To explore the structure of information represented in speech and language embeddings, we used t-SNE to project the high-dimensional embedding spaces down to two-dimensional manifolds for visualization. This projection was computed separately for the speech embeddings (from the encoder network) and the language embeddings (from the decoder network). Each data point in the scatter plots (Fig. 2) corresponds to a speech or language embedding for a unique word. For each unique word (n = 13,347) we averaged the embeddings across instances throughout the transcript to get one embedding per word. We replicated the analysis using the first instance or random instances of each word and obtained similar results. We then applied t-SNE to the averaged embeddings with perplexity = 50. To better understand the structure of this two-dimensional space, we colored the data points (corresponding to word embeddings) according to several speech and language features: phonemes, place of articulation (PoA), manner of articulation (MoA), and part of speech (PoS). Phonemes, PoA, and MoA capture acoustic and articulatory features of speech, whereas PoS captures lexical categories. We obtained phoneme classes from the Carnegie Mellon Pronouncing Dictionary ^59^, which provides 39 classes (37 in our dataset). We further classified the phonemes based on their place of articulation (total 9 classes, 9 in our dataset) and manner of articulation (total 9 classes, 9 in our dataset) according to the general american english consonants of the International Phonetic Alphabet. Because each word consists of multiple phonemes, we took the first phoneme for each word. We replicated the following visualizations and classification analyses for the second, third, and fourth phonemes of each word separately and obtained similar results. To extract part of speech information, we used the part of the speech tagging process available in the NLTK python package (total 12 classes, 11 in our dataset). We removed classes with less than 100 occurrences (less than 1% of the data, resulting in 27 phoneme classes, 9 PoA classes, 9 MoA classes, and 5 PoS classes).

### Classification of speech and linguistic features

To quantify the information encoded in the embeddings, we trained multinomial logistic regression classifiers (using the L2 penalty and default *C* = 1.0 in scikit-learn) to predict phonetic (phonemes, PoA, MoA) and lexical categories (PoS) separately for both speech and language embeddings. We used a ten-fold cross-validation procedure with temporally-contiguous training/test folds to train and evaluate classifier performance. On each fold of the cross-validation procedure, embeddings were standardized and reduced to 50 dimensions using PCA. To establish a baseline for comparing classifier accuracy, we trained dummy classifiers that learned to predict the most frequent class. Since the distribution of classes in our dataset was unbalanced, we used the balanced accuracy metric to evaluate classification performance ^60^. Balanced accuracy is calculated as the proportion of correct predictions per class averaged across all classes. This results in a value between 0 and 1, with higher values indicating better classification performance. For instance, a random classifier that always predicts the most frequent class will result in a balanced accuracy of 1 divided by the number of classes, which is at the chance level. Essentially, the balanced accuracy metric assesses how well the classifier can differentiate between different classes while minimizing misclassifications due to unbalanced data.

### Preprocessing the ECoG recordings

The ECoG preprocessing pipeline was intended to mitigate artifacts due to movement, faulty electrodes, line noise, abnormal physiological signals (e.g., epileptic discharges), eye blinks, and cardiac activity ^61^. We built a semi-automated analysis pipeline to identify and remove corrupted data segments (e.g., due to epileptic seizures or loose wires) and mitigate other noise sources using FFT, ICA, and de-spiking methods ^62^. We then bandpassed the neural signals using a broadband (75–200 Hz) filter and computed the power envelope, a proxy for each electrode’s average local neural firing rate ^63^. The signal was z-scored and smoothed with a 50 ms Hamming kernel. 3000 samples were trimmed at each end of the signal to avoid edge effects. Signal preprocessing was performed using custom preprocessing scripts in MATLAB 2019a (MathWorks).

### Electrode-wise encoding models

To map the Whisper embeddings onto the neural activity, we used linear regression to estimate encoding models for each electrode and lag relative to word onset. To construct the outcome variable, we averaged the neural signal across a 200 ms window at each lag (at 25 ms increments) for each electrode across all words. Using a ten-fold cross-validation procedure, we trained two sets of encoding models to predict the wordy-by-word neural signal magnitude based on either speech or language embeddings. Within each training fold, we standardized the embeddings and used PCA to reduce the embeddings to 50 dimensions. We then estimated the regression weights using ordinary least-squares multiple linear regression from the training set and applied those weights to predict the neural responses for the test set. To assess model performance, we calculated the Pearson correlation between the predicted neural signal and the actual neural signal for each held-out test fold. The correlations were average across the ten test folds. This procedure was repeated at 161 lags from -2,000 to 2,000 ms in 25-ms increments relative to word onset; the same predictor embeddings were used at each lag.

### Electrode selection

To identify significant electrodes, we used a randomization procedure. At each iteration, we performed a random shift in the assigned embeddings to each predicted signal, thus disconnecting the relationship between the words and the brain signal while preserving the order between the different embeddings. The random shift was restricted to avoid rolling the assignment inside the context window. We then performed the entire encoding procedure for each electrode on the mismatching words. We repeated this process 1,000 times. After each iteration, the encoding model’s score was calculated based on the maximal value minus the minimal value across all 161 lags for each electrode. For each patient, we then took the maximum value for each permutation across all electrodes. This resulted in a distribution of 1,000 maximum values for each patient, which was used to determine the significance of all electrodes. For each electrode, a *p*-value was computed as the percentile of the original maximum-minimum values of the encoding model across all lags from the null distribution of 1,000 similarly calculated values. Performing a significance test using this randomization procedure evaluates the null hypothesis that there is no systematic relationship between the brain signal and the corresponding word embedding. This procedure yielded a family-wise error rate corrected *p-*value for each electrode, correcting for the multiple lags ^64^. Electrodes with *p*-values less than 0.01 were considered significant.

### Differences in the overall magnitude of encoding performance

To identify electrodes with significant differences in the magnitude of encoding performance for speech and language embeddings, we used the same randomization procedure described in the electrode selection section. We only statistically evaluated differences in model performance for electrodes with significant encoding performance for at least one model (see “Electrode selection” above). For each permutation, we computed the difference in model performance by subtracting the two maximal encoding performance (correlation) values for each electrode across all 161 lags. This resulted in a distribution of 1,000 difference values between the encoding performance for speech embeddings and language embeddings at each electrode. For each electrode, a *p*-value was computed as the percentile of the non-permuted maximum difference values between encoding performance for speech and language embeddings across all lags from the null distribution of 1,000 difference values. To correct for testing across multiple electrodes, we used FDR correction ^65^. Electrodes with *q*-values less than 0.005 (significance of 0.01 standardized for the two-sided test) were considered significant differences in model performance. We used the same procedure to identify electrodes that showed a significant difference in the magnitude of encoding performance between speech production and comprehension.

### Differences in lag-by-lag encoding performance

To test for significant differences in electrode-wise encoding performance between the encoding performance for the speech and language embeddings for each lag, we used a paired-sample permutation procedure: in each permutation, we randomly shuffled the labels of all observations for both models (we obtained a correlation coefficient for each fold during a 10-fold validation procedure, thus collecting 10 observations per electrode for each model). Then, we computed the difference between encoding performance for speech and language embeddings. We compute the exact null distribution of different values for the 10 observations (2^10^ = 1,024 permutations). For each lag, a *p*-value was computed as the percentile of the non-permuted difference relative to the null distribution of 1,024 difference values. To correct for multiple lags, we used the FDR correction procedure ^65^. Lags with *q*-values less than 0.025 (significance of 0.05 for the two-sided test) were considered to be significant.

We used a similar procedure to test for significant differences in electrode-wise encoding performance for the speech and language embeddings averaged across electrodes in different ROIs: we randomly shuffled the labels of all observations (10 × *n*, where 10 is the number of folds and *n* corresponds to the number of electrodes in the ROI) and computed the difference between mean encoding performance for the speech embeddings and language embeddings. This process was repeated 10,000 times, resulting in a distribution of 10,000 difference values. For each lag, a *p*-value was computed as the percentile of the non-permuted difference relative to the null distribution, and FDR correction was applied to correct for multiple lags. Lags with *q*-values less than 0.025 (significance of 0.05 for the two-sided test) were considered to be statistically significant.

### Differences in the temporal lag of peak encoding performance

To test for significant differences in the temporal dynamics of encoding performance between ROIs, we performed independent-samples *t*-tests. First, we hypothesized that for production, peak encoding in electrodes in IFG would occur significantly earlier than in electrodes in somatomotor (SM) and auditory (STG) areas. To test this hypothesis, we performed an independent-samples *t*-test (one-sided) on the lags at peak encoding for electrodes in the given ROIs. Second, we hypothesized that for comprehension, peak encoding in electrodes in IFG would occur significantly later than peak encoding in electrodes in SM and STG. We performed an independent-samples *t*-test (one-sided) on the lags at peak encoding for electrodes in the given ROIs to test this hypothesis. To test whether the peak encoding performance for electrodes in a given ROI occurred significantly before or after word onset, we performed one-sample *t*-tests (two-sided) on the lags at peak encoding for electrodes in the given ROI against lag 0 (word onset).

## Supplementary Information

**Supp. Fig. 1.**
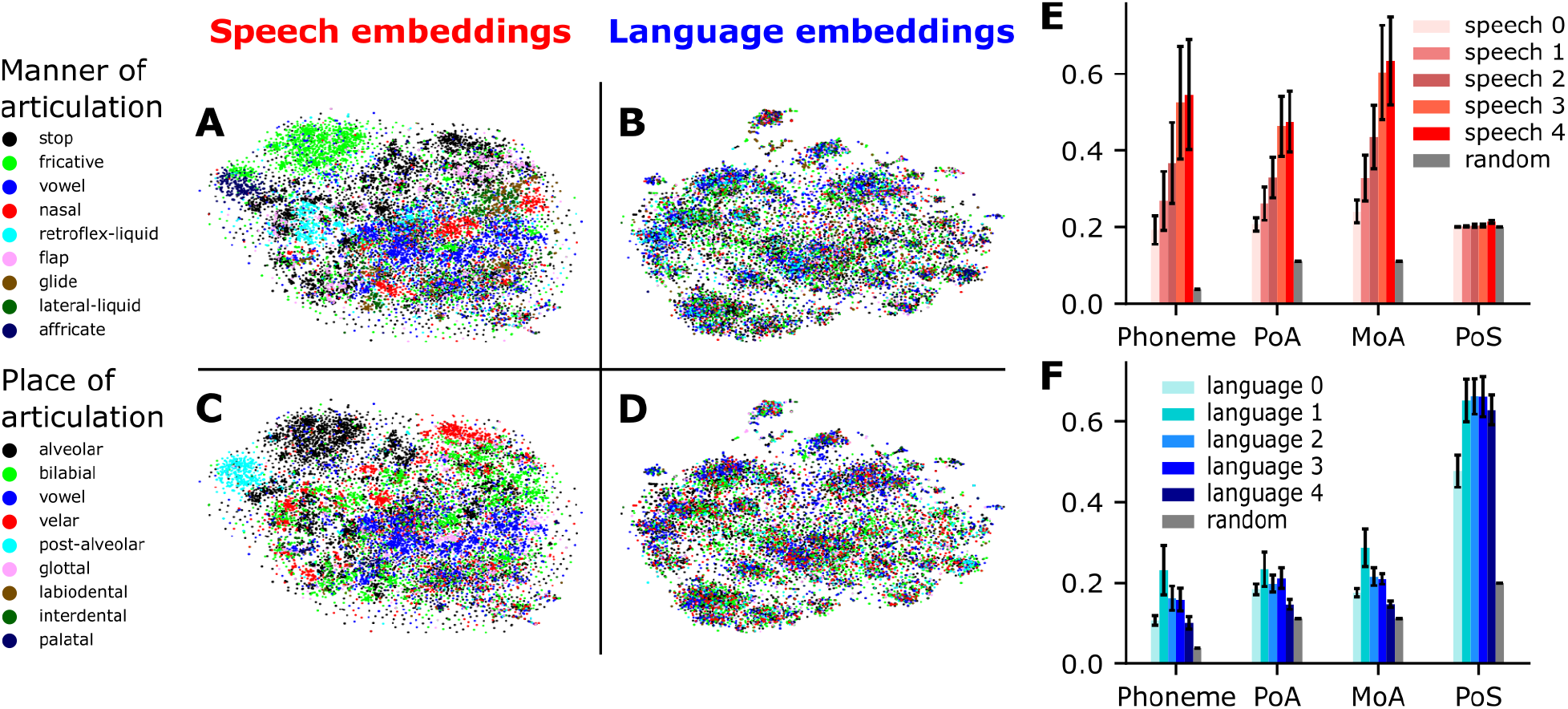
Representations of phonetic and lexical information in Whisper. (A-D) Visualization of speech embeddings and language embeddings in a two-dimensional space estimated using t-SNE. Each data point corresponds to the embedding for either an audio segment (speech model) or a word token (language model) for a unique word (averaged across all instances of one word). Clustering according to phonetic categories (manner of articulation, MoA; and place of articulation, PoA) is visible in speech embeddings (A, C) but not in language embeddings (B, D). (E, F) Classification of phonetic information (phoneme, MoA, PoA) and lexical information (PoS) based on embeddings taken from the different layers of the whisper encoder and decoder network. The last layer of speech embeddings (speech 4) shows best classification accuracy for phonetic information and relatively low classification accuracy for lexical information (E). The third layer of language embeddings (language 3) show best classification accuracy for lexical information and relatively low classification accuracy for phonetic information (F).

**Supp. Fig. 2.**
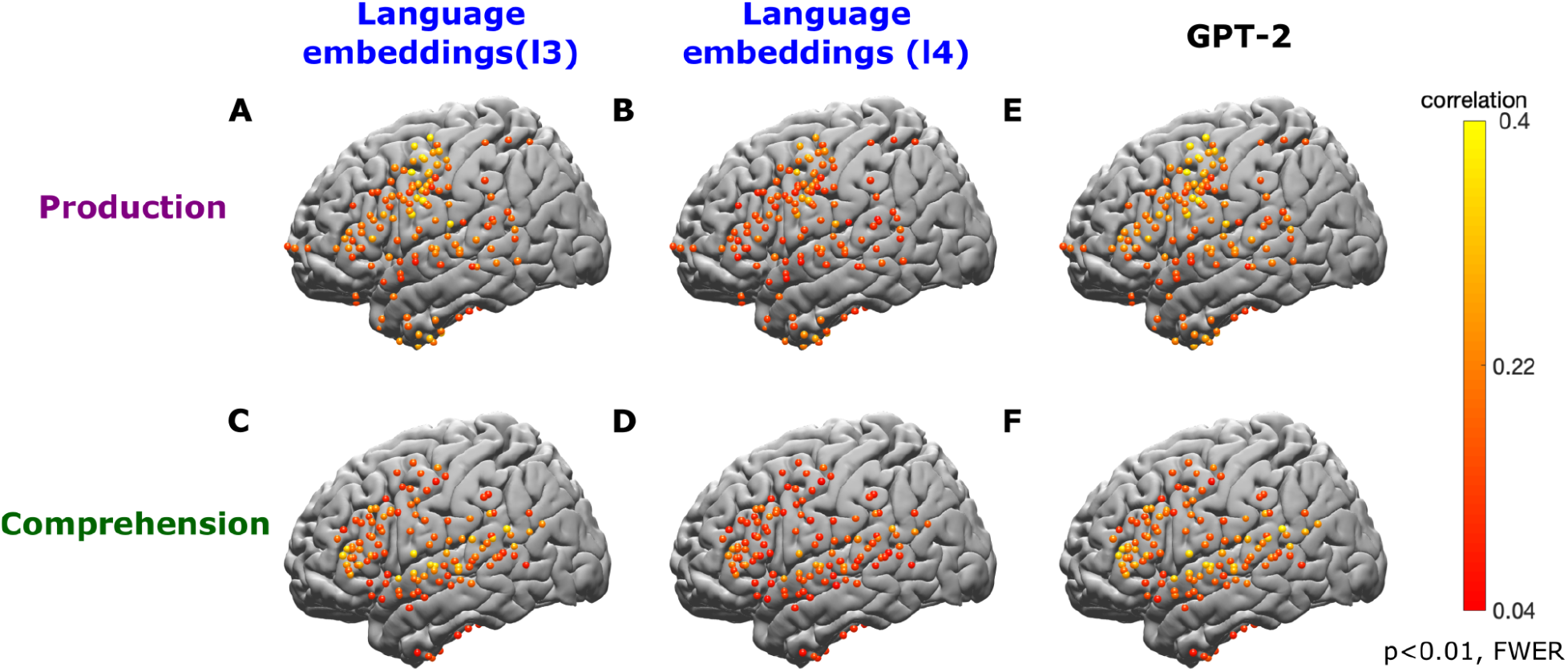
Comparing language embeddings across layers and models. We compared encoding performance for language embeddings from Whisper decoder layer 3 and 4 to encoding performance for language embeddings from GPT-2 layer 8. Encoding performance is shown for significant electrodes during speech production and comprehension, corrected for multiple comparisons. The color indicates the maximum correlation value across lags per electrode.

**Supp. Fig. 3.**
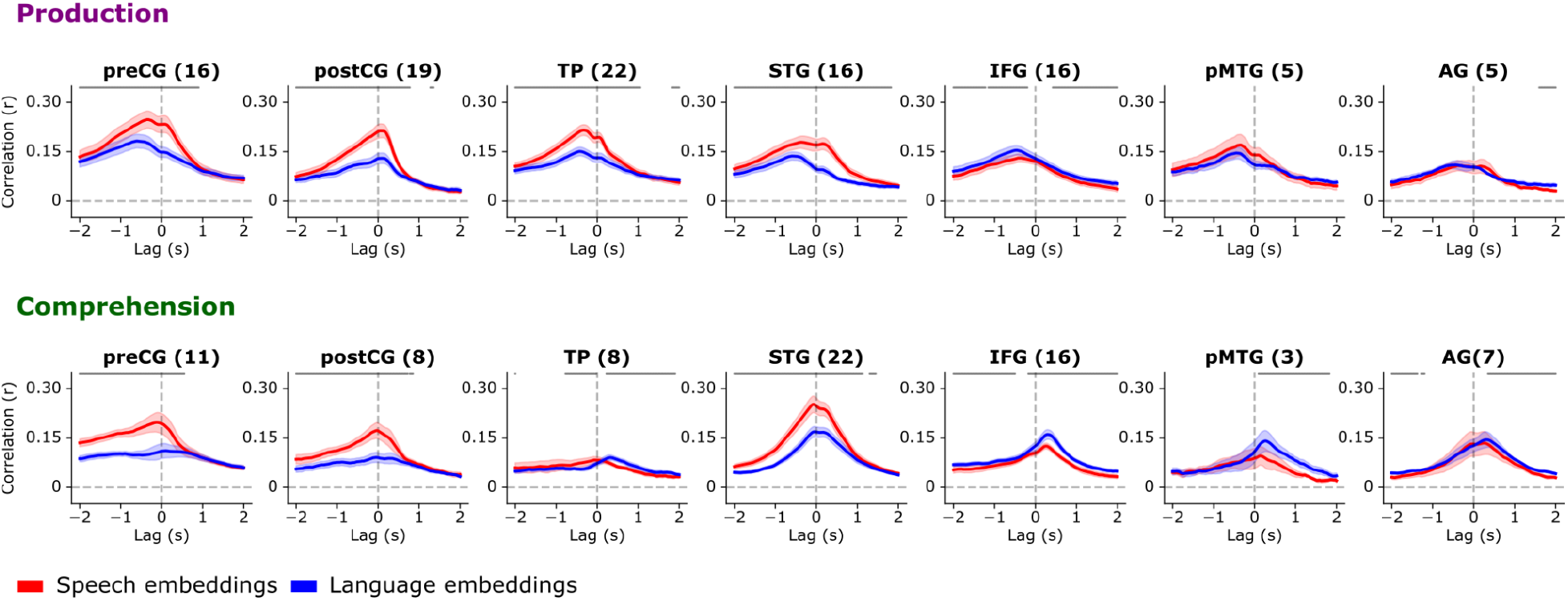
Average speech and language encoding for ROIs. Electrode-wise encoding performance values were averaged per ROI for speech (red) and language (blue) embeddings. For each ROI, encoding performance was averaged across electrodes that showed a significant difference in maximum encoding performance (across lags) between speech and language embeddings (number of electrodes in parentheses). The shaded color represents the standard error. Asterisks indicate a lag-wise significant difference in ROI-wise encoding performance between speech and language embeddings (*q* < 0.01, FDR corrected).

**Supp. Fig. 4.**
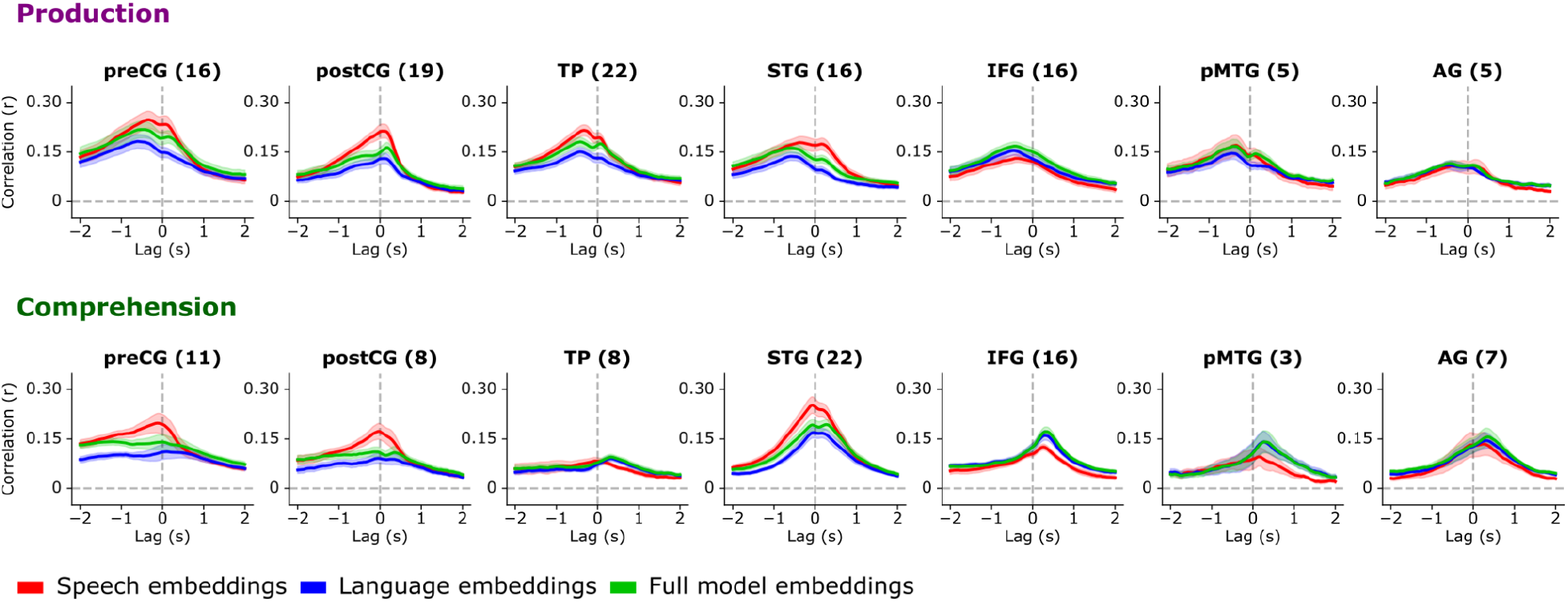
Average encoding for the full Whisper model. Comparing electrode-wise encoding performance averaged per ROI using speech embeddings (red), language embeddings (blue), and “full” model embeddings (green). Full model embeddings were extracted from layer 3 of the Whisper decoder network while keeping the connection to the encoder network intact. The full model, therefore, includes both speech and language information.

**Supp. Fig. 5.**
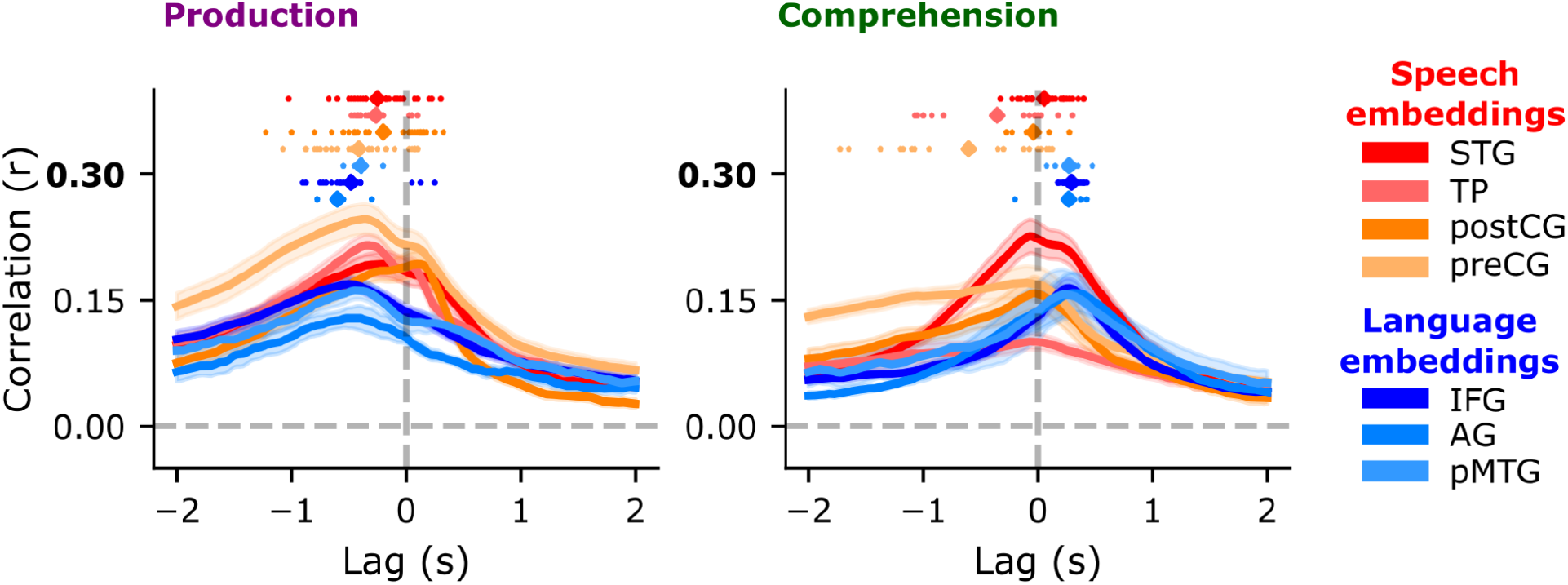
Temporal dynamics of speech production and speech comprehension across different ROIs. The best-performing model was selected for each ROI to assess lags. Colored dots show the encoding peak for each electrode per ROI. To assess whether the peak occurred significantly before or after word onset, we performed a one-sample *t*-test comparing the sample mean of encoding peaks per ROI against 0. We show that for speech production the encoding peak occurs before word onset in all ROIs (AG (M = -600 ms, SD = 196 ms, t(3) = -5.3, p < 0.05), IFG (M = -480 ms, SD = 278 ms, t(22) = -8.08, p < 0.001), pMTG (M = -395 ms, SD = 114 ms, t(4) =-6.9, p < 0.005), STG (M = -250 ms, SD = 301 ms, t(26) = -4.24, p < 0.001), TP (M = -264 ms, SD = 150 ms, t(26) = -9.11, p < 0.001), postCG (M = -199 ms, SD = 383 ms, t(29) = -2.79, p < 0.01), preCG (M = -414ms, SD = 362 ms, t(28) = -6.06, p < 0.001). For speech comprehension, encoding peaked after word onset in high-level language areas (AG (M = 268 ms, SD = 186 ms, t(7) = 3.81, p < 0.01), pMTG (M = 271 ms, SD = 72 ms, t(28) = 21.4, p < 0.005), IFG (M = 292 ms, SD = 72 ms, t(28) = 21.4, p < 0.001). In sensory areas, STG (M = 54 ms, SD = 186 ms, t(32) = 1.65, p > 0.05) and postCG (M = -38 ms, SD = 139 ms, t(10) = -0.87, p > 0.05) the peak occurred around word onset. In preCG (M = -603ms, SD = 624ms, t(18) = -4.1, p < 0.001) and TP (M =-355ms, SD = 503ms, t(12) = -2.44, p < 0.05), the peak occurred before word onset.

**Supp. Fig. 6.**
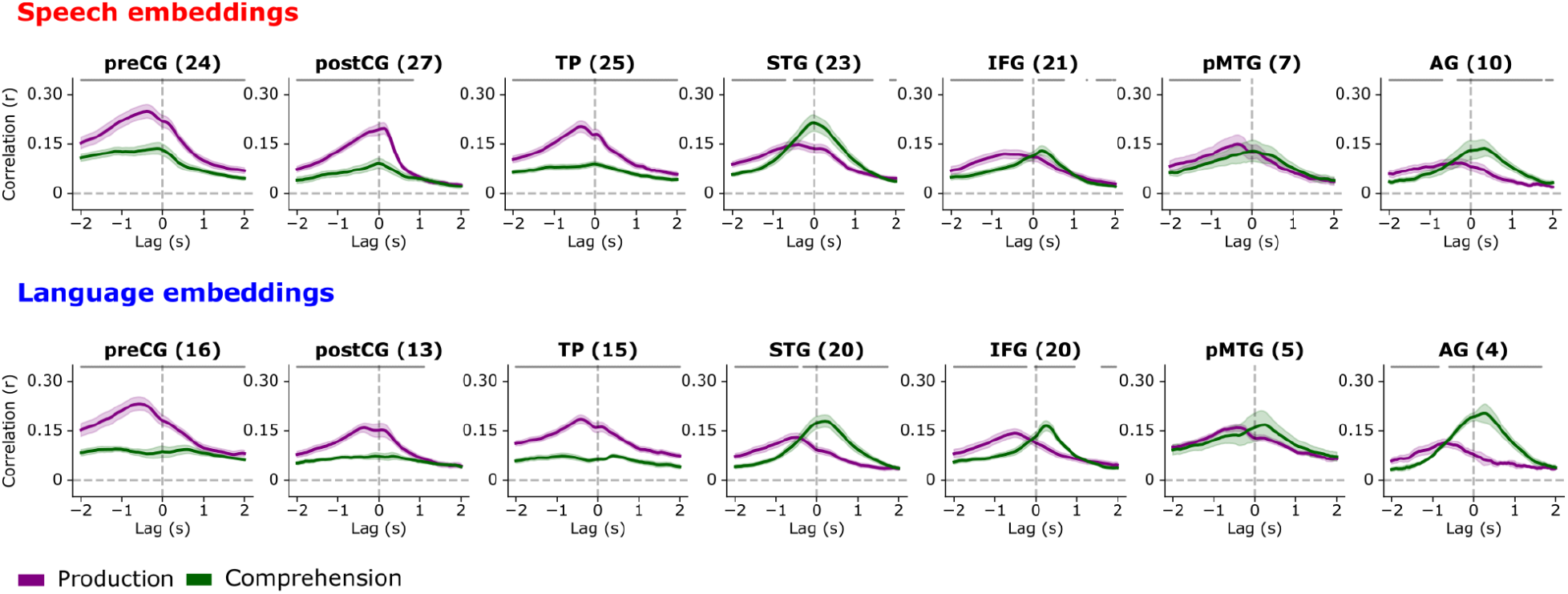
Average ROI-level encoding for speech production and comprehension. Electrode-wise encoding performance values were averaged per ROI for production (purple) and comprehension (green). For each ROI, encoding performance was averaged across electrodes that showed a significant difference in maximum encoding performance (across lags) between production and comprehension (number of electrodes in parentheses). The shaded color represents the standard error. Asterisks indicate a lag-wise significant difference in ROI-wise encoding performance between production and comprehension (*q* < 0.01, FDR corrected).

**Supp. Table 1.** Patient demographics and clinical characteristics.

| Participant | 1 | 2 | 3 | 4 |
| --- | --- | --- | --- | --- |
| Age | 53 | 26 | 48 | 24 |
| Sex | F | M | F | M |
| Number of electrodes implanted | 104 | 125 | 255 | 192 |
| Hours of speech recorded | 17 | 37 | 17 | 29 |
| Number of words recorded | 79654 | 213473 | 117800 | 109282 |
| Number of comprehension words | 47642 | 109967 | 71754 | 60608 |
| Number of production words | 32012 | 103506 | 46046 | 48674 |
| Neuropsychological testing scores | VCI: 105<br>POI: 107<br>PSI: 120<br>WMI: 102 | VCI: 89<br>POI: 75<br>PSI: 65<br>WMI: 74 | VCI: 107<br>POI: 104<br>PSI: 111<br>WMI: 114 | VCI: 145<br>POI: 96<br>PSI: 86<br>WMI: 95 |
| Pathology/epilepsy type/seizure focus | posterior temporal lobe (neocortical) epilepsy. IEEG results were consistent with a seizure focus adjacent to or overlying the known posterior temporal cortical lesion | left anteromedial temporal lobe epilepsy | Right anteromedial temporal lobe epilepsy. ICEEG localized ictal onsets to the right temporal pole and right hippocampus | Focal epilepsy arising from the left hemisphere with a broad focus involving the left temporal neocortex (superior, middle, inferior temporal gyri) left frontal operculum, inferior postcentral gyrus, insula |
| Implant | Left grid, strips, and depth | Left grid, strips, and depth | Bilateral strips and depths, and a left grid | Left grid, strips, and depth |

